# Mutation rate estimate and population genomic analysis reveals decline of koalas prior to human arrival

**DOI:** 10.1101/2025.05.15.654135

**Authors:** Toby G. L. Kovacs, Nicole M. Foley, Luke W. Silver, Elspeth A. McLennan, William J. Murphy, Carolyn J. Hogg, Simon Y. W. Ho

## Abstract

The koala (*Phascolarctos cinereus*), an iconic Australian marsupial, has experienced substantial historical and contemporary population declines. Identifying the drivers of these declines has been hindered by limited genomic data and by uncertainty regarding the koala mutation rate. Here, we provide the first direct estimate of the koala mutation rate, based on four parent-offspring trios, yielding a mean of 6.12×10□□ mutations per base pair per generation (95% confidence interval: 5.03–7.45×10□□). Using this estimate of the rate, we reconstructed the demographic history of koalas using 458 whole-genome sequences sampled across their entire range. Our results refine the estimated timing of past changes in population size, suggesting a large decline beginning ∼100 kya, before the arrival of modern humans in Australia. The koala population then split into five genetic populations 6–30 kya, which are now spread across >3500 km on the east coast of Australia. We also use the koala mutation rate to infer recombination maps for each population, confirming lower recombination rates in marsupials than in eutherian mammals. These findings provide critical insights into the evolutionary history of koalas, while highlighting the impact of using species-specific evolutionary rates in the inference of demographic histories and recombination landscapes. Our estimates of the genome-wide mutation rate and population-specific recombination maps for koalas provide valuable resources for future evolutionary and conservation analyses of marsupials.

## Introduction

The Australian continent contains a rich diversity of endemic flora and fauna, which endured severe environmental shifts throughout the late Cenozoic. The continent was dominated by wet forests during the Paleogene but underwent drastic changes during the Miocene as the Australian tectonic plate drifted northwards (White 1994). Widespread aridification was marked by the retraction of rainforests to the coasts, the opening of forests, and the expansion of grasslands and deserts (Byrne et al. 2011). Intense glacial and interglacial climate cycles during the Plio-Pleistocene then drove further shifts towards arid and fire-prone environments (White 1994). These changes have underpinned the evolution of Australian biota, driving adaptation to arid environments, the isolation of taxa in remnant mesic habitats, and the extinction of many lineages, especially rainforest specialists (Byrne 2008; Byrne et al. 2008; Byrne et al. 2011). More recently, the first arrival of modern humans 65–47 kya was followed by the extinction of Australia’s megafaunal species, changes to fire frequency and habitats (O’Connell and Allen 2015; Clarkson et al. 2017).

Marsupials are among the most recognizable components of the Australian fauna. These include the koala (*Phascolarctos cinereus*), which is the sole extant member of the family Phascolarctidae. The koala lineage diverged from its closest extant living relatives, the wombats, ∼36 Mya (Duchêne et al. 2018), but fossil evidence suggests that the late Oligocene and early Miocene contained a large diversity of koala species across multiple genera (Black et al. 2014; Black 2016; Crichton et al. 2023). The modern koala species appeared at least 350 kya, while most of the other species went extinct by the end of the Pleistocene (Black et al. 2014). Bioclimate modelling has estimated that koala habitat on the east coast experienced substantial contractions during the Last Glacial Maximum before expanding to its present-day range during the current interglacial period (Adams-Hosking et al. 2011). Contemporary koala populations inhabit an enormous range (>3500 km) but have experienced recent declines as a result of land clearing, disease, hunting, feral dog attacks, vehicle strikes, and bushfires (Menkhorst 2008; Adams-Hosking et al. 2016; Cristescu et al. 2023), leading to their listing as Endangered in Queensland (QLD), New South Wales (NSW), and the Australian Capital Territory (ACT) in 2022.

Genetic studies have suggested historically small population sizes in koalas, with museum samples from the 1800s containing low mitochondrial diversity (Tsangaras et al. 2012). Studies of mitochondrial DNA have suggested that population sizes were stable over the last 50,000 years, before expanding after the Last Glacial Maximum (Neaves et al. 2016). Although evidence from exons has pointed to pronounced bottlenecks >300 kya (Lott et al. 2022), whole-genome analyses have suggested that koala populations declined precipitously after modern humans arrived on the continent (Johnson et al. 2018; De Cahsan et al. 2025). This aligns with global patterns of anthropogenic impacts on megafauna, including in Australia (Johnson 2016; Van Der Kaars et al. 2017; Adesanya et al. 2023). However, the demographic inferences from koala genomes relied on mutation rate estimates from distantly related eutherian mammals (humans and mice), introducing uncertainty in the timing of population-size changes. Therefore, a comprehensive analysis using a more relevant mutation rate can provide a clearer picture of the patterns and drivers of demographic change in koalas.

Mutation rates can be estimated using three main methods: (i) the phylogenetic method, which infers the mutation rate from an estimate of the substitution rate; (ii) the mutation-accumulation method, which requires the sampling of a population for many generations; and (iii) the parent-offspring trio method (Pfeifer 2020). The phylogenetic method has historically been the most widely used, given the relative ease of estimating the substitution rate using molecular-dating methods with fossil calibrations. However, the substitution rate is expected to underestimate the mutation rate, because a proportion of *de novo* mutations are removed by purifying selection over time (Ho et al. 2011; Yoder and Tiley 2021). Mutation-accumulation and trio-based methods tend to yield broadly similar estimates of mutation rates (Wang and Obbard 2023), but the former method is only feasible for model organisms with short generations.

There has been rapid growth in mutation rate estimates from parent-offspring trios, particularly for vertebrate species (Bergeron et al. 2023; Wang and Obbard 2023). Although many estimates have been obtained for eutherian mammals, there have only been two estimates for marsupial species, each based on a single parent-offspring trio. The only estimate from an Australian marsupial (Tasmanian devil, *Sarcophilus harrisii*) was based on individuals from a highly inbred captive population and might not reflect the mutation rate in wild populations (Bergeron et al. 2023). Therefore, obtaining further estimates of mutation rates in Australian marsupials, particularly for the speciose order Diprotodontia, has the potential to produce substantial improvements in demographic and evolutionary inferences.

Another benefit of obtaining a species-specific mutation rate is accurately inferring other evolutionary parameters, such as recombination maps. Until recently, producing recombination maps was a laborious process that involved genotyping individuals of known pedigree through generations to allow the identification of crossover events between chromosomes (Burbrink et al. 2025). This barrier has now been overcome with the availability of reference-quality long-read assemblies for species of conservation concern (R. N. Johnson et al. 2018) and machine-learning approaches that can infer recombination maps from population genomic data (Adrion et al. 2020). These methods have been used to estimate the recombination landscape for a number of non-model species (Bredemeyer et al. 2023; Foley et al. 2024), but their accuracy has been hindered by uncertainty in mutation rates (Adrion et al. 2020).

Recombination rates in marsupials are expected to be lower than those of eutherian mammals, based on their linkage map length (Dumont and Payseur 2008). However, an accurate genomic estimation of the recombination landscape using a relevant mutation rate is required to confirm this. Species-specific recombination maps can also be used to assess whether genes of conservation concern are in regions of low or high recombination. This determines the impact of linked selection (Nordborg et al. 1996) and the likelihood of successfully incorporating diversity through translocations, helping conservation managers better tailor management strategies to a population’s needs.

Here we estimate the genomic mutation rate of koalas from four parent-offspring trios and use it to infer demographic history from a large set of whole-genome sequences. We further leverage this mutation rate estimate to generate population-specific recombination maps for koalas and demonstrate their use by querying the recombination context of immune genes. Our study shows that the koala population began to decline prior to the arrival of modern humans on the Australian continent.

## Results

### Estimation of the koala mutation rate

We sequenced the genomes of 12 koalas from three families, comprising seven parents and five offspring, with an average coverage of 32× across the samples. The inferred relationships in one trio differed from that recorded in the studbook and so this trio could not be used to estimate mutation rates. We identified 100 unique *de novo* mutations in the remaining four offspring after filtering for false discovery following the methods used by Bergeron et al. (2023). We estimated a mean mutation rate of 6.12×10^-9^ mutations/bp/generation (95% confidence interval: 5.03–7.45×10^-9^) across the four trios (Fig. 1*a*), using the false-negative rate (0.068) and the size of the callable genome (average 68.6% total genome size) (supplementary table S1). The mean estimate can also be expressed as 8.74×10^-10^ mutations/bp/year, assuming a generation length of 7 years (Phillips 2000). The detected mutations had a transition to transversion ratio of 2.23 (Fig. 1*b*).

**FIG. 1.**
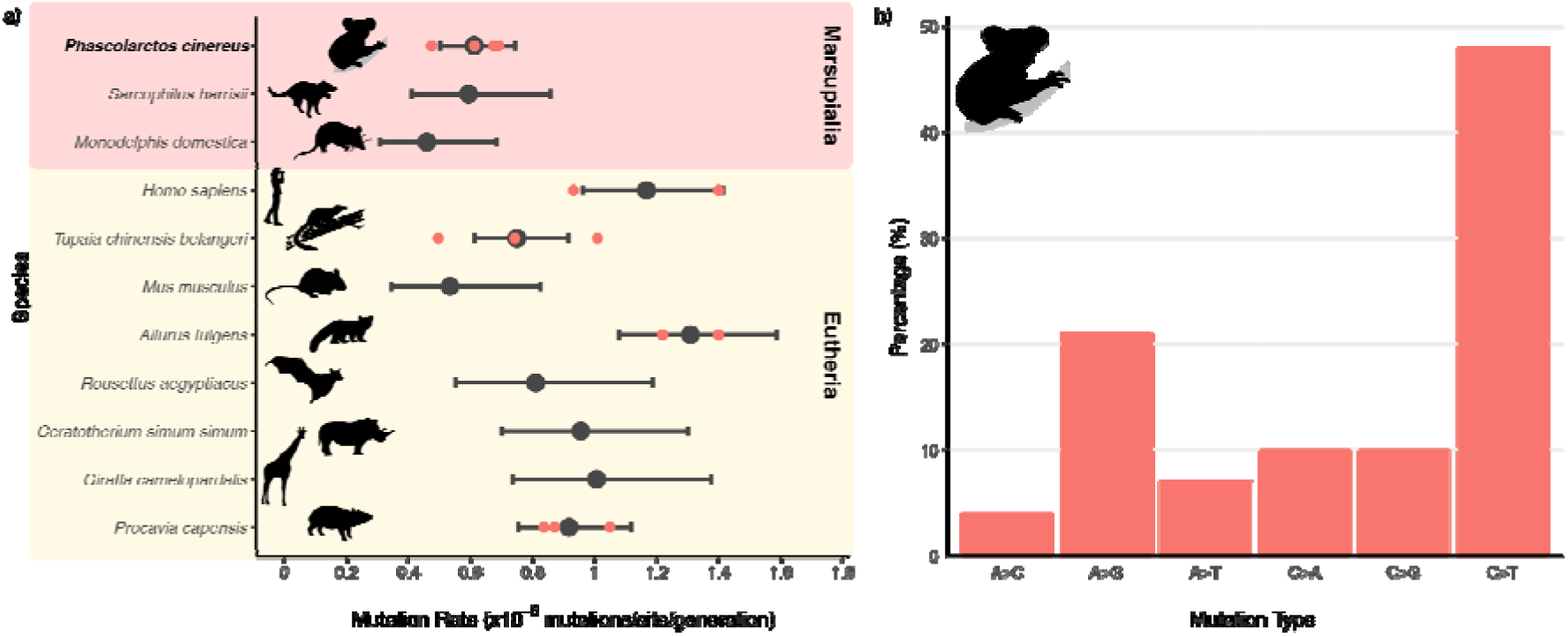
Germline mutation rate of the koala and other mammals. (*a*) Germline mutation rates inferred in the koala (*Phascolarctos cinereus*) based on four parent-offspring trios. Individual estimates for each trio are shown in orange. The average mutation rate is shown in black and 95% confidence intervals are indicated with error bars. Other mutation rate estimates are from Bergeron et al. (2023) and can be found in supplementary table S2. Silhouettes were sourced from Phylopic (http://phylopic.org). (*b*) Mutation spectrum for koala genomes. The percentage of each mutation type is shown after collapsing reverse complements.

### Estimation of demographic history

We performed population demographic analyses using 458 genomes of wild koalas, including all 413 samples from the Koala Genome Survey (Hogg et al. 2023; McLennan et al. 2024) and 45 additional samples from Kangaroo Island and the Mount Lofty Ranges (South Australia) to fill geographical gaps. Our phylogenetic network of wild and captive koalas showed a weakly structured population, with samples broadly grouping into the five populations that were recognized in a previous study (McLennan et al. 2024): North Queensland (N QLD), South-East Queensland/North New South Wales (SE QLD/ N NSW), Middle New South Wales (M NSW), Southern New South Wales (S NSW), and Victoria (VIC) (Fig. 2*b*). We also show that most of the additional South Australian koalas form a nested clade within the Victorian population. However, a few koalas from the Mount Lofty Ranges were spread throughout the Victorian clade, suggesting that multiple lineages could be present in South Australia.

**FIG. 2.**
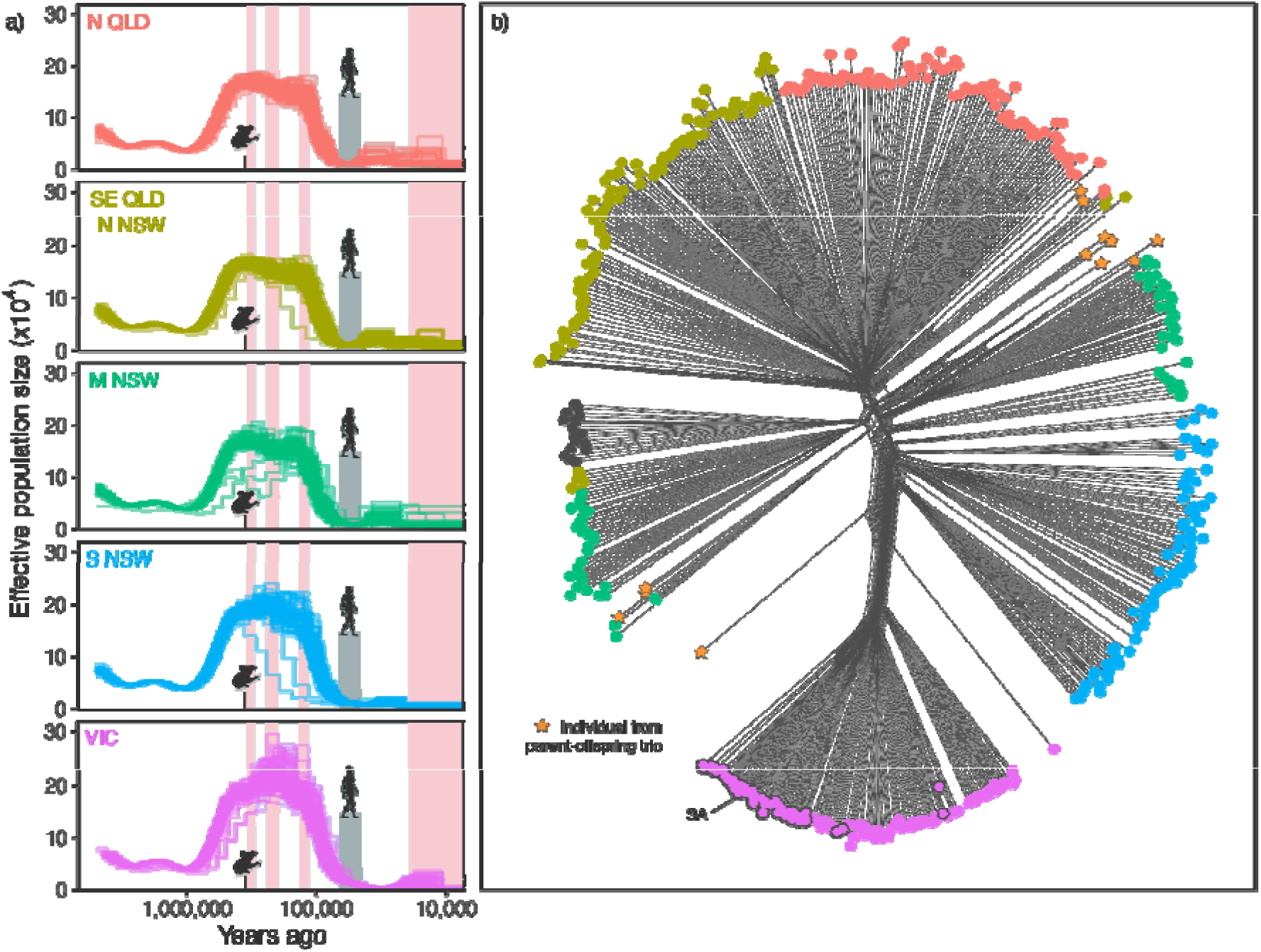
(*a*) Historical effective population sizes estimated using PSMC for 458 individuals across the five genetic populations. Backgrounds indicate the interglacial (warmer and wetter; pink) and glacial (cooler and drier; blue) periods of the last three glacial cycles. The dark grey column and human silhouette indicate the period in which humans are believed to have arrived and spread across Australia (65–47 kya; O’Connell & Allen 2015; Clarkson et al. 2017). The koala silhouette indicates the earliest known fossil of the modern koala species (Black et al. 2014). (*b*) Phylogenetic network of all 474 koala samples based on Euclidean genetic distances. Colours correspond to the five genetic clusters shown in (*a*). Individuals that form parent-offspring trios used for mutation-rate estimation are indicated by yellow stars. Koalas from Narrandera, which represent a mix of two populations, are shown in dark grey. Grey outlines highlight koalas from South Australia (SA), which are nested within the Victorian population.

We estimated the historical demography of the koala using the pairwise sequentially Markovian coalescent (PSMC; Li and Durbin 2011). Our results suggest that koala populations expanded 700 kya and remained stable until ∼120 kya, when they experienced severe and prolonged reductions to reach their minimum population size ∼60 kya (Fig. 2*a*). This pattern showed little variation within and between populations and did not appear to be influenced by the level of sequencing coverage (supplementary fig. S1). Our results also suggest that population sizes remained small following this bottleneck, although PSMC is not particularly reliable for inferring demographic trends on recent timeframes (Patton et al. 2019).

When we used the default time intervals designed for analysing human genomic data, several of our samples produced PSMC plots with recent spikes in population size (supplementary fig. S2). These spikes are known artefacts in PSMC analyses and previous studies have shown that they can be removed by subdividing recent time intervals (Hilgers et al. 2025).

We also found that, in cases where similar artefacts were produced when grouping the first four intervals (denoted as “4+…”), subdividing the first time interval into two or four intervals (as “2+2+…” and “1+1+1+1+…”) sometimes removed these peaks (supplementary fig. S2). In some cases, however, when grouping the first four intervals did not introduce an artefact, subdividing the first interval introduced an artificial peak in population size, suggesting that this issue should be assessed on a case-by-case basis.

Demographic inference using the sequentially Markovian coalescent can be improved by the addition of multiple samples per population, with SMC++ considered the state-of-the-art method for analysing samples of more than eight genomes (Terhorst et al. 2017). Our estimates of historical population sizes using SMC++ showed increased resolution, with a larger number of shorter time intervals, and with greater similarity in demographic plots between populations when increasing the number of knots (supplementary fig. S3). Our plots using 50 knots showed a similar pattern to that observed in our PSMC results, with koala populations increasing more gradually from 700 to 200 kya before declining between 110 and 40 kya to ∼10% of their maximum population size (Fig. 3*a*). Following the bottleneck, the three northern populations (N QLD, SE QLD/N NSW, and M NSW) expanded at 15 kya before stabilizing. The South New South Wales population recovered slightly, whereas the Victorian population does not show any signs of recovery.

**FIG. 3.**
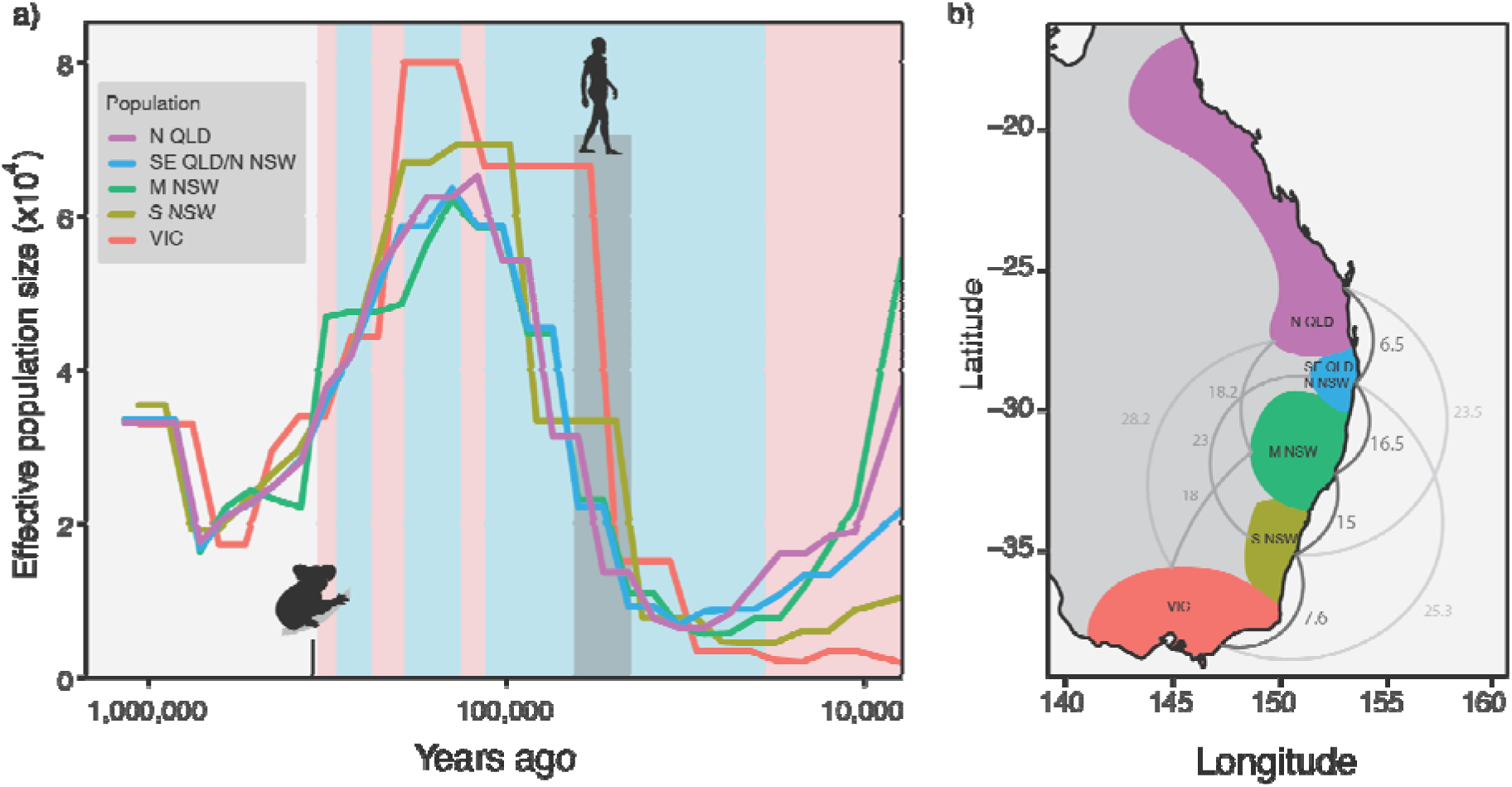
(*a*) Historical population sizes estimated using SMC++ for each of the five genetic groups of koalas. Pink and blue backgrounds indicate the interglacial (warmer and wetter) and glacial (cooler and drier) periods of the last three glacial cycles. The dark grey column and human silhouette indicate the period in which humans are believed to have arrived and spread across Australia. The koala silhouette indicates the earliest known fossil of the modern koala species (Black et al. 2014). (*b*) Population split times (kya) estimated using SMC++ for all pairs of populations. The split times are darker and larger between more closely located populations.

We also used SMC++ to estimate the split times between pairs of koala populations. Divergences among the populations were estimated to have occurred in the last 30 ky, following a general isolation-by-distance pattern (Fig. 3*b*; supplementary fig. S4). This suggests that the severe bottleneck occurred in the ancestral population of all extant koalas, with modern lineages subsequently diverging as the population sizes recovered. Based on the pairwise estimates between geographically adjacent populations, we found that the ancestral koala population split at 15−16.5 kya to form the three ancestral populations of i) N QLD and SE QLD / N NSW, ii) M NSW, and iii) S NSW and VIC. The two northernmost and two southernmost populations were the last to split, diverging from the remaining populations at 6.5 and 7.6 kya, respectively.

### Estimation of the population specific koala recombination maps

We estimated differing recombination rates between populations using ReLERNN (Adrion et al. 2020) (Fig. 4), with the VIC population having the highest mean recombination rate of 1.40×10^-9^ crossovers/bp/generation (Q1−Q3: 0.83–1.84×10^-9^ crossovers/bp/generation) and the SE QLD/N NSW population with the lowest recombination rate 0.74×10^-9^ crossovers/bp/generation (Q1−Q3: 0.51–0.91×10^-9^ crossovers/bp/generation) (Fig. 4; supplementary table S3). The VIC population had a significant difference in recombination rate compared to the four other populations (Mann-Whitney test, all comparisons *p* < 2.2×10^-16^) (Fig. 4). The recombination landscape was highly correlated when using either the pan-mammal or koala-specific mutation rate (M NSW rho=0.74, p < 2.2×10^-16^; N QLD rho=0.65, p < 2.2×10^-16^; VIC rho=0.66, p < 2.2×10^-16^). However, the mean recombination rate was significantly higher when using the pan-mammal compared with the koala-specific mutation rate for the three populations analysed (Mann-Whitney test, all comparisons *p* < 2.2×10^-16^).

**FIG. 4.**
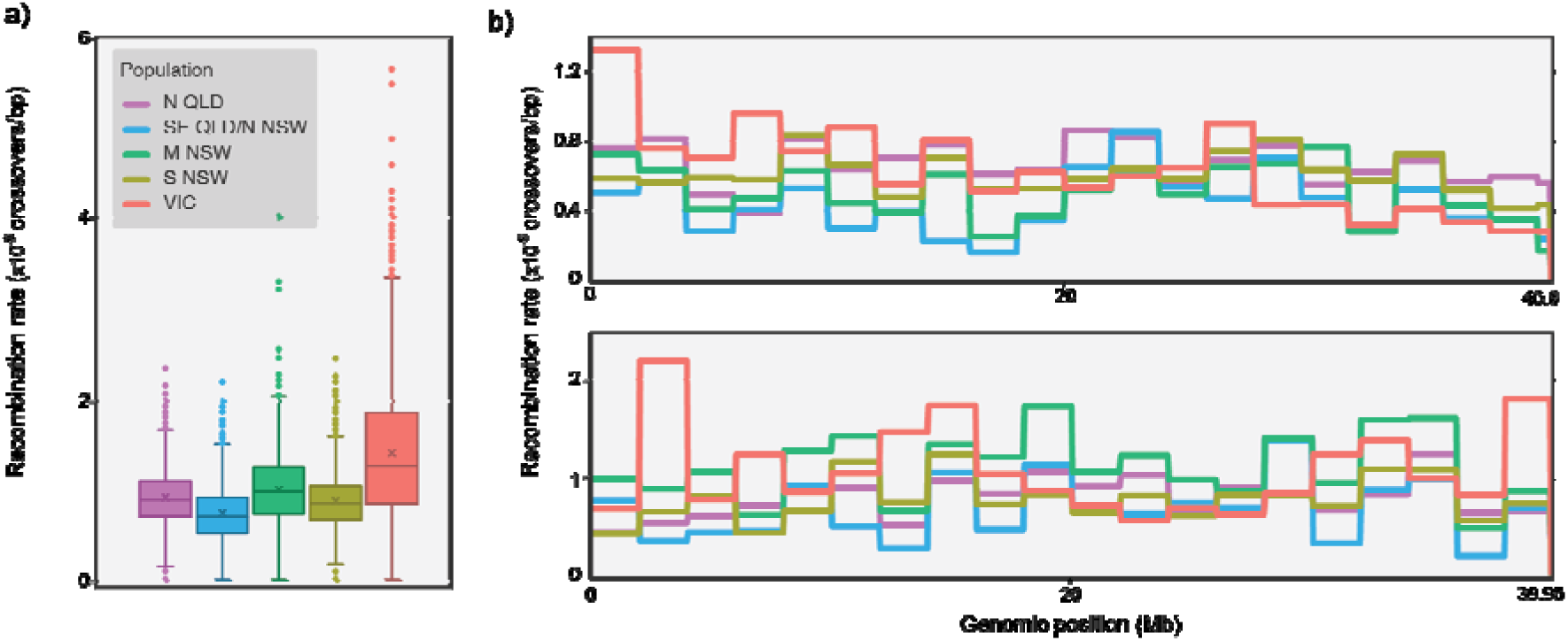
Variation in recombination rates across koala populations. (*a*) Boxplots showing genome-wide recombination rates. In each box, the × and the horizontal line represent the mean and the median recombination rate, respectively. Whiskers extend to the most extreme data points within 1.5 times the interquartile range from the first and third quartiles. Points beyond this are plotted. (*b*) Variation in recombination rates among populations across the two largest scaffolds.

The components of the koala immune repertoire varied considerably in terms of their recombination context (supplementary table S4). The tissue-resident memory T cells were consistently found in high-recombining genomic regions across all populations. In contrast, major histocompatibility complex (MHC) class III genes were generally found in lower-recombining regions across all populations (supplementary table S4). However, the recombination context of some components differed between populations. In the VIC population, the MHC extended class II components occur in low-recombining genomic regions (1st decile), while the same components are found in predominantly high-recombining regions (9th decile) in the population from SE QLD/N NSW.

## DISCUSSION

### Germline mutation rate in the koala

Our sequencing and analysis of koala genomes from parent-offspring trios has yielded a mutation rate estimate of 6.12×10^-9^ mutations/bp/generation (95% confidence interval: 5.03– 7.45×10^-9^), similar to those reported for the Tasmanian devil (5.95×10^-9^ mutations/bp/generation) and Virginia opossum (*Didelphis virginiana*; 4.60×10^-9^ mutations/bp/generation) but considerably lower than that inferred for modern humans (11.6×10^-9^ mutations/bp/generation) (Bergeron et al. 2023). The koala yearly mutation rate (0.87×10^-9^ mutations/bp/year) is lower than those of the Tasmanian devil (1.27×10^-9^ mutations/bp/year) and Virginia opossum (2.35×10^-9^ mutations/bp/year), due to the shorter generations of these two other marsupials (Lewin and Eyre-Walker 2025). The yearly koala rate is comparable to those of eutherian mammals with similar generation times but higher than that of humans (0.43×10^-9^ mutations/bp/year; Lewin and Eyre-Walker 2025). Our estimate of the koala mutation rate is the first for Diprotodontia, the largest extant marsupial order, providing a valuable resource for evolutionary and demographic analyses of this group.

We obtained similar estimates of mutation rates from genomes sequenced from museum skins and from fresh blood. This result demonstrates that older tissue samples can provide a useful genomic resource with strict site filtering, despite theoretical concerns (Bergeron et al. 2022). Although we sequenced to a lower depth than Bergeron et al. (2023), we found similar percentages of the genome as being callable after the same strict filtering. However, our study has also revealed potential risks in relying on studbook records for parental identifications. We also detected high relatedness between the parents in one trio, suggesting that their founding individuals were likely to be related. This led to a small number of sites for which the parents were homozygous for different alleles, meaning that this trio contributed minimally to the calculation of the false-negative rate.

Compared with previous estimates of mutation rates in marsupials, which only used a single trio each (Bergeron et al. 2023), our analysis of four replicate trios has enabled us to more accurately estimate the distribution of mutations across classes. The distribution across mutation classes for koalas was broadly similar to that observed in primates (Fig. 1*b*; Versoza et al. 2025), despite >120 My since the divergence between marsupial and eutherian mammals (Álvarez-Carretero et al. 2022) and mutation spectra diverging between some eutherian mammals (Beichman et al. 2023).

Our study provides an estimate of a key evolutionary parameter that can enhance genomic analyses of koalas and other marsupials. Further estimates of mutation rates from a broader range of marsupial species will allow us to test whether the drivers of mutation rates are consistent with those seen in eutherian mammals. Previous studies have reported relationships between the number of *de novo* mutations and both maternal and paternal age at conception (Jónsson et al. 2017), as well as a paternal bias in mutation contributions (Manuel et al. 2022). Marsupials could differ in the rate at which mutations accumulate because they give birth after a short gestation period, with most development occurring in the pouch rather than *in utero* (Smith & Keyte 2020). Teasing apart how *de novo* mutations arise during this process could help to reveal the broader determinants of mutation rates. Among species, mutation rate variation is correlated with generation time (Bergeron et al. 2023; Zhu et al. 2025), effective population size (Lynch et al. 2016; Zhu et al. 2025), and genome size (Lynch et al. 2016). It is unclear whether these patterns hold in marsupials, but their contrasting developmental strategies make them an interesting test case for evolutionary theories of mutation rate variation in mammals.

### Population declines coincide with glacial environmental shifts

Our results provide a compelling illustration of the impact of the mutation rate used to scale the timing of demographic inferences. Notably, our application of the koala mutation rate led to demographic plots that were drastically different from those scaled using the human mutation rate. Although our PSMC curves have the same shape as those previously inferred for koalas, we found that population declines occurred prior to the arrival of modern humans ∼65–47 kya (O’Connell and Allen 2015; Clarkson et al. 2017), whereas previous studies suggested that these declines followed human arrival (Johnson et al. 2018; De Cahsan et al. 2025). There has been much debate over when humans first reached the continent (O’Connell et al. 2018; Bradshaw et al. 2023), but the arrival of humans into southern Australia, and the majority of the koala’s range, most likely occurred towards the more recent end of this scale. While both our PSMC and SMC++ results show that the koala population started to decline prior to human arrival, our PSMC estimates suggest sharp bottlenecks while our SMC++ estimates suggest a more gradual decline that partially overlaps with the presence of humans. In addition, compared with previous studies (Johnson et al. 2018; De Cahsan et al. 2025), our PSMC and SMC++ analyses both infer an earlier expansion of koala populations 700–200 kya, aligning with the oldest known fossil of the present-day koala 350 kya (Black et al. 2014; Fig. 2*a*, Fig. 3*a*).

We estimated that a bottleneck occurred in the ancestral koala population before it split into the present-day populations in the last 30 ky. These results are substantially different from previous reports of the populations diverging 250–200 kya (De Cahsan et al. 2025) or the ancestor diverging into three populations >300 kya (Lott et al. 2022). After the populations began to diverge, we estimated slight recoveries in the three northernmost populations but not in the two southern populations. This pattern is consistent with the present-day diversity in each of the populations, pointing to the influence of pre-European processes. However, it is well documented that southern populations experienced more severe recent bottlenecks due to a prolific fur trade around the turn of the 20th century (Phillips 1990; Menkhorst 2008), leading to decreased diversity and increased inbreeding (McLennan et al. 2024). The impact of these recent processes on SMC++ analyses is not well understood, and they could potentially be driving the observed patterns in historical sizes of the southern populations.

This bias could also explain the reduced resolution (number of time intervals) in the analysis of the Victorian population, for which a slightly later bottleneck was inferred than for the other populations. Given that the Victorian population was not isolated until after the bottleneck, the inferred later bottleneck is unlikely to be biologically meaningful.

Population declines during the last 110 ky coincide with the onset of the most recent glacial period. During this period, koala ranges became increasingly restricted to habitat refugia along the eastern and southwest coasts (Adams-Hosking et al. 2011). This is likely to have seen the final separation of the eastern koala populations from the now-extinct western population through the emergence of the Nullarbor (meaning ‘no tree’) Plain, the treeless southern central region of Australia. During the mid-Pleistocene, the Nullarbor Plain harboured a diverse collection of herbivores that included arboreal species such as koalas and tree kangaroos, suggesting a more diverse flora than at present (Prideaux et al. 2007). This would have provided a connection between eastern and western eucalypt forests, before the landscape transitioned to semi-arid shrubland by ∼70 kya to form a vast biogeographical barrier between the western and eastern koala populations (Van Der Kaars et al. 2017; De Deckker et al. 2021; Adesanya et al. 2023). The appearance of this barrier would have led to a substantial reduction in the effective population size of koalas, matching the declines observed in our demographic models.

Koalas would have become increasingly restricted to the east and west coast as they approached the Last Glacial Maximum (Adams-Hosking et al. 2011). This is consistent with the declines inferred from SMC++, which suggest a continued decline until ∼20 kya. Modern humans arrived on the Australian continent after the koala population had already started to decline, but humans have been credited with the extinction of the last remaining Western Australian population (Balme et al. 1978; Prideaux et al. 2010). Population expansions and divergences after the Last Glacial Maximum also coincide with the expansion of habitat across the east coast of Australia to the koala’s present-day range of more than 3500 km (Adams-Hosking et al. 2011).

Historical population sizes estimated using the sequentially Markovian coalescent (e.g., PSMC and SMC++) rely on the assumption that a population’s inverse instantaneous coalescent rate is directly correlated with its effective population size. However, a reduction in the inverse instantaneous coalescent rate can be caused not only by a reduction in population size, but also by increasing structure with reduced migration and gene flow (Mazet et al. 2016; Chikhi et al. 2018; Mather et al. 2020). Therefore, it is possible that the inferences from these methods indicate a decline in connectivity between koala populations during this period instead of reductions in size. However, decreased population size and increased population structure would both support the notion that koala habitat became more restricted during the last glacial period. It is likely that these two processes occurred in tandem.

### Population-specific koala recombination landscape

Our inferred recombination maps contained mean recombination rates ranging from 0.74×10^-9^ to 1.40×10^-9^ crossovers/bp/generation, an order of magnitude lower than the average rate in animals of 25.2 ×10^-9^ crossovers/bp/generation (Stapley et al. 2017). However, the estimated recombination rates in koalas are more consistent with the low rates inferred for other marsupials from linkage maps, such as 2.3×10^-9^ crossovers/bp/generation in the gray short-tailed opossum (*Monodelphis domestica*) (Samollow et al. 2007, Samollow et al. 2010) and 5.1×10^-9^ crossovers/bp/generation in the tammar wallaby (*Notamacropus eugenii*) (given a genome length of 3.3 Gb; Wang et al. 2011). Reduced recombination rates in marsupials have been explained by fewer double-stranded breaks during meiosis (Marín-Gual et al. 2022), but the evolutionary drivers remain unknown.

When recombination is reduced, the Hill-Robertson effect reduces the effectiveness of selection and the effective population size at linked sites, increasing genetic drift and lowering local levels of diversity (Charlesworth et al., 1993; Nordborg et al., 1996). Therefore, lower recombination rates are expected to further threaten genetically depleted species. We inferred differing recombination rates across the five koala populations, with the highest rate in the VIC population. This result suggests that although the VIC population has the lowest level of genetic diversity due to a recent bottleneck (McLennan et al. 2024), its high rate of genome-wide recombination means that it is likely to experience lower levels of linked selection. This could potentially explain how this population has successfully increased in population size following its near extinction in the early 20th century (Whisson and Ashman 2020).

Population-specific variation in recombination rates has previously been found in a number of species. In humans and other animals, recombination has been found to be driven by variation in the *PRDM9* alleles which determine the genomic locations of recombination hotspots (Paigen and Petkov 2018). Other causes of population-specific recombination rates include variation in inversions, chromatin structure, and epigenetics (Stapley et al. 2017). Our recombination maps provide a useful resource for determining whether the same drivers of recombination rates occur in marsupials.

Producing a recombination map for a heavily managed threatened species could help to predict the success of mixing genetic diversity through translocation programs, one of the primary objectives of genomics-based conservation management. In sexually reproducing populations, meiotic recombination (along with the random assortment of chromosomes) creates new combinations of alleles in the offspring as homologous chromosomes cross over and exchange genetic material (Burbrink et al. 2025). Traditional conservation management strategies like translocations are most likely to create new combinations of genetic variants in high-recombining parts of the genome, where exchange occurs most frequently. However, if the goal of a translocation is to reduce or eliminate deleterious variation in a gene that resides in a low-recombining genomic region, it will be harder to remove these variants without also losing diversity at linked sites. Our recombination maps have highlighted how much the recombination context can vary between genes and how this pattern can vary across different populations. Our results suggest that translocations of koalas between populations are more likely to generate novel combinations of alleles in tissue-resident memory T (TRM) cell genes compared with MHC class III genes. With greater knowledge about the specific genes or variants conferring resistance and susceptibility to *Chlamydia* infection in koalas, these resolved recombination maps will help conservation managers to predict whether translocation experiments are likely to affect genomic regions of interest.

Previous estimates of recombination maps using ReLERNN in mammal species for which estimates of mutation rates are unavailable (Bredemeyer et al. 2023; Foley et al. 2024) have used a previously inferred ‘pan-mammal’ rate (Kumar and Subramanian 2002). Although this approach enables robust comparison of the recombination landscape within a species, it does not allow direct comparison with other species. Our results confirm the original assessment that ReLERNN produces correlated recombination landscapes for differing mutation rates, but applying an overestimate of the mutation rate, as is the case when applying the ‘pan-mammal’ rate to koalas, results in overestimation of the recombination rates (Adrion et al. 2020). These outcomes highlight another benefit of producing species-specific mutation rates for a diverse range of organisms.

Although our analysis was based on a high-quality, long-read-based genome assembly, it also highlighted some insufficiencies in this genome assembly strategy for the study of the immune loci. Immune genes are innately difficult to assemble because they are highly heterozygous, often evolve through gene duplication, and contain complex structural rearrangements (Logsdon et al. 2025). Although long-read sequencing technologies have evolved to account for repetitive genomic regions, they still struggle to assemble highly heterozygous parts of diploid genome assemblies (Rice et al. 2020). This was reflected in the fact that some immune loci were assigned to scaffolds that were too small for their rate of recombination to be inferred with confidence. This was an issue for genes within the immunoglobulin kappa locus and genes located within the immunoglobulin heavy-chain locus (supplementary table S4). Our results highlight the need for complete telomere-to-telomere haploid genomic sequences, not only for humans, apes, and agricultural and companion animals (Nurk et al. 2022; Bredemeyer et al. 2023; Rice et al. 2020), but also for species of conservation concern.

## Conclusions

Genome-scale analyses of demographic and evolutionary history are allowing increasingly fine-grained inferences of complex historical processes. We have shown, however, that our ability to do so in non-model organisms, which often contain the most fascinating histories, is hindered by gaps in our knowledge of fundamental evolutionary parameters. With a growing catalogue of mutation rate estimates for organisms across the Tree of Life, our demographic modelling will become increasingly accurate and will allow us to resolve the drivers of historical population-size dynamics.

In the koala, our use of a direct estimate of the mutation rate led to a shift in the estimated timing of population-size changes, with important consequences for identifying the drivers of population declines. In particular, we found that late Pleistocene population contractions occurred prior to the arrival of humans on the continent. This shifts the narrative away from anthropogenic declines of koalas and highlights the substantial ecosystem changes that have occurred during the history of the continent. Broader analyses of the Australian fauna are required to determine whether this pattern is unique to koalas, or if ecosystem changes have driven similar declines in other species.

We have also demonstrated that using species-specific mutation rates allows more accurate estimation of recombination maps. Using these maps, we characterized the recombination context of koala immune genes, identifying genes that are more likely to be affected by linked selection and more likely to benefit from translocations. These results provide a useful context to help conservation managers tailor management strategies to a population’s needs.

## Materials and Methods

### Whole-genome resequencing of parent-offspring trios

Our data set comprised 12 resequenced genomes from five male and seven female koalas, from five putative parent-offspring trios from three families. Eight individuals forming three trios were sourced from skins at the Australian Museum and from a captive-bred colony at Featherdale Sydney Wildlife Park. Four live individuals forming two trios (two full siblings) were selected from the Taronga Conservation Society Australia’s breeding program, with DNA extracted from blood. All genome sequences were produced in the first stage of the Koala Genome Survey (Hogg et al. 2023; McLennan et al. 2024). We downloaded bam files from the Amazon Web Services Open Data Sponsorship program (Australasian Genomes; https://koalagenomes.s3.ap-southeast-2.amazonaws.com/index.html). Details of DNA extraction, sequencing, and read mapping were reported previously (Hogg et al. 2023; McLennan et al. 2024). The average genome coverage across koala samples forming trios was 31×. *De novo* mutations, false-positive rates, and false-negative rates were calculated using the methods described by Bergeron et al. (2022), with the additional false-positive steps introduced by Bergeron et al. (2023).

The Koala Genome Survey variant call format (VCF) file was updated to include all available koala genomes, including captive-bred individuals, as well as wild representatives from South Australian populations on Kangaroo Island (Gates et al. 2025) and the Mount Lofty Ranges. We used individual genomic variant call format (GVCF) files for each sample as input to Illumina’s DRAGEN Germline Pipeline (v4.3.6; Illumina), using the koala reference genome (GCA_002099425.1_phaCin_unsw_v4.1; Johnson et al. 2018) to perform joint genotyping across all samples. This uses the DRAGEN gVCF PopGen Genotyper (4.3.6; Illumina) to produce a single, multi-sample variant calling file (VCF) used for all downstream analyses. Depth (DP) was calculated for all sites in all samples using bcftools (v1.17), by summing the localized allelic depths (LAD). Our unfiltered VCF contained 59,694,393 SNPs. Variants were filtered using bcftools to remove sites that did not pass Illumina’s QUAL filter and had read depth < or <60, leaving 47,655,085 SNPs. Finally, we filtered out variants that were missing in more than one individual, leaving 17,573,439 SNPs. Custom scripts based on those developed in Bergeron et al. (2022) and Bergeron et al. (2023) for the estimation of mutation rates are available at https://github.com/tobykovacs796/koalamutrate.

### Phylogenetic network

The filtered VCF was read into a genlight object in R using *read.PLINK*. Euclidean distances were calculated using *bitwise.dist* in the package poppr, with the “scale_missing” setting.

Using these genetic distances, we inferred a phylogenetic network using SplitsTree 6 (Huson and Bryant 2024). Individuals were then assigned to populations based on their maximum ancestry in previous *fast*STRUCTURE analyses (McLennan et al. 2024). Custom scripts for demographic analyses are available at https://github.com/tobykovacs796/koalamutrate.

### Demographic inference

Historical population sizes were estimated using a suite of methods that use the sequentially Markovian coalescent (SMC) as well as the site frequency spectrum. SMC methods are generally most accurate between 1000 and 100,000 generations ago (Patton et al. 2019), with PSMC being more accurate for deeper timeframes in this range, and newer methods, such as SMC++, being more accurate for more recent timeframes due to the consideration of the site frequency spectrum (Terhorst et al. 2017). In order to demonstrate the impact of the mutation rate on demographic inference, we ran analyses using PSMC and SMC++.

We analysed all 458 genomes from wild koalas using PSMC (Li and Durbin 2011). We trimmed the koala reference genome (GCA_002099425.1_phaCin_unsw_v4.1; Johnson et al. 2018) using vcftools (Danecek et al. 2011) to remove all contigs <50 kb and those putatively corresponding to the X chromosome (Hogg et al. 2024). We generated the input file for PSMC (Li and Durbin 2011) by calling variants for each individual using the bam file and trimmed reference genome using samtools mpileup (v1.6) and bcftools (v1.3.1), following the instructions given for PSMC (Li and Durbin 2011). PSMC was run using the time intervals “4+25×2+4+6”, assuming a generation length of 7 years (Phillips 2000), and with our estimate of the per-generation mutation rate. Individual demographic plots were grouped according to the previously assigned genetic clusters of koalas (McLennan et al. 2024).

Because some of our initial PSMC plots displayed population spikes in single recent time intervals, we tried splitting the earlier time intervals as done by Hilgers et al. (2025) (2+2+25×2+4+6 and 1+1+1+1+25×2+4+6). This removed the spikes in some plots but added them to other plots where they did not originally occur. Hence, the final plot was selected to minimize these extreme, single-interval peaks, which are a known artefact of these methods.

Historical population sizes were also estimated for each of the five genetic populations using SMC++. Koalas were assigned to a population based on their highest ancestry as identified previously (McLennan et al. 2024). Koalas from Narrandera were not included in any population for SMC++ analyses because they are the result of recent translocations from SE QLD / N NSW and VIC populations (McLennan et al. 2024). Captively bred koalas from Dubbo, which were not included in previous analyses of population structure (McLennan et al. 2024), were placed across M NSW and S NSW based on our phylogenetic network.

Because these koalas are likely to contain recently mixed ancestry, they were excluded from SMC++ analyses. Koalas from South Australia, which were nested within the Victorian population in our phylogenetic network, originated from serial bottlenecking and inbreeding and so were not included in the analysis of Victorian koalas using SMC++.

For our demographic analyses using SMC++ (Terhorst et al. 2017), we used vcftools (Danecek et al. 2011) to filter the VCF to keep only biallelic sites and remove all contigs <50 kb and those that have been putatively identified as forming the X chromosome (Hogg et al. 2024). Historical population sizes were inferred in SMC++ using *estimate*, calculating the composite likelihood across several distinguished lineages. Distinguished lineages were selected based on the largest group of unrelated individuals that could be found in each population, with up to 10 distinguished lineages. We ran SMC++ with default settings with our estimated mutation rate, across the time interval 10^3^–10^6^ years. Initial analyses using the default number of knots (8) showed limited resolution over key timeframes of interest, so analyses were rerun with the number of knots set to 16, 32, and 50. These results showed increasing resolution with the number of knots (supplementary fig. S3), by introducing more inflection points and inferring different effective population sizes for a larger number of shorter time intervals. Several runs with 32 or 50 knots failed shortly after starting, with the error: *erroneous average coalescent time*. These runs were restarted until they completed successfully. The timing of population separation was inferred in SMC++ (Terhorst et al. 2017) using the function *split* and the marginal estimates for each population when using 50 knots. Custom scripts for demographic analyses are available at https://github.com/tobykovacs796/koalamutrate.

### Recombination maps

A subset of unrelated individuals from each of the five populations were selected to infer population-specific recombination landscapes. Relatedness (kinship) between individuals was estimated within each of the five populations identified in McLennan et al. (2024) using vcftools *--relatedness2*, which implements the KING method (Manichaikul et al. 2010). We wrote a script to randomly select individuals and iteratively build a group of unrelated individuals containing five females and five males. The algorithm began by randomly choosing one individual, then repeatedly searching the population for an unrelated individual, adding each as it was found, until either a group of ten was assembled or no additional unrelated individuals could be added. When multiple valid sets of ten were possible, we prioritized individuals to maximize genome coverage and ancestry representation. Variants for these individuals were subsetted from the unfiltered multi-sample VCF. ReLERNN requires that SNPs residing in transposable elements be masked before the analysis. As such, we identified the genome-wide transposable element content of the reference genome using RepeatMasker (version Open 3.2.6 A.F.A. Smit, R. Hubley & P. Green RepeatMasker at http://repeatmasker.org). This program was run using the -qq parameter and using the Zoonomia repeat library (Osmanski et al. 2023). To prepare variants for generating a recombination map, we removed variants overlapping a transposable element annotation in the reference genome. We further filtered variants, removing variants within 5 bp of an indel and those that did not meet the following quality criteria:e’%QUAL<30 | INFO/DP<16 | INFO/DP>62 | QD<2 | FS>60 | SOR>10 | ReadPosRankSum <-8 | MQRankSum <-12.5 | MQ<40’ in bcftools (https://github.com/samtools/bcftools). Using VCFtools (https://vcftools.github.io/man_latest.html), we removed indels and subset the data to include only biallelic SNPs for each clade for further analysis. Custom scripts used to contextualize and average the raw ReLERNN output are available at https://github.com/eutherialab/Foley_XLRD.

To model the genome-wide recombination rate for each koala population, we used ReLERNN (https://github.com/kr-colab/ReLERNN), a deep learning approach that uses recurrent neural networks (Adrion, Galloway, and Kern 2020). Our estimate of the mutation rate was incorporated into the analysis. ReLERNN was run using the simulate, train, predict, and bscorrect modules with default settings. Given that we were interested in broadly evaluating the differences in recombination rates among populations, we averaged rates in 2 Mb blocks with a 50 kb step. A non-parametric Mann-Whitney test was used to determine if there was a significant difference between the populations with the highest and lowest inferred recombination rates.

We then investigated whether there was a difference between recombination rates inferred using an empirically estimated mutation rate for koalas compared with a ‘pan-mammal’ yearly mutation rate of 2.2×10^9^, estimated in a previous analysis of branch lengths (Kumar and Subramanian 2002). This was multiplied by 7 to produce a per-generation mutation rate for koalas to allow direct comparison with our empirically estimated mutation rate. We reran ReLERNN for a subset of populations using the pan-mammal mutation rate. Tests for normality were conducted using qq-norm in R. Spearman’s rank correlation test was used to evaluate differences in the inferred recombination landscapes. A Mann-Whitney test was used to determine if there were differences in the mean of the rates between recombination maps.

### Recombination context of koala immune genes

Per-population, raw recombination rates were ordered from lowest to highest and assigned to deciles, where the 1st and 10th deciles represent the lowest and highest recombination rates, respectively. Previous analyses have identified and annotated immune gene clusters in the koala genome (R. N. Johnson et al. 2018). These included comprehensive annotation of the major histocompatibility complex, T-cell receptors, immunoglobulins, cytokines (R. N. Johnson et al. 2018), additional MHC genes (Silver et al. 2022; Silver et al. 2024), toll-like receptors (Cian et al. 2024), and natural killer cells (Morris et al. 2015). For each gene, we list the recombination decile that corresponds to those genomic coordinates for each of the five koala populations.

## Data availability

Raw whole genome re-sequences are available in The National Center for Biotechnology Information (NCBI) under BioProject PRJNA940526 at https://www.ncbi.nlm. nih.gov/bioproject/PRJNA940526. All outputs including the position of *de novo* mutations and relatedness values are available at https://github.com/tobykovacs796/koalamutrate and the supplementary material.

## Acknowledgements

We thank Taronga Zoo for taking blood samples from their captive koalas (Animal Ethics Committee approval 4a/06/22) and the Australian Museum and Featherdale Sydney Wildlife Park for providing skins, and all institutions for access to the studbook. We thank Eilish McMaster for help with data visualization, and Zhiliang Chen and the Illumina team for assistance with DRAGEN.

